# In silico spatial transcriptomic editing at single-cell resolution

**DOI:** 10.1101/2023.08.25.554778

**Authors:** Jiqing Wu, Viktor H. Koelzer

## Abstract

**Motivation:** Generative artificial intelligence (AI) has enabled fundamental breakthroughs in visual content creation by text-guided editing. However, the utility of such generative models remains largely understudied for processing increasingly complex bioimage data.

**Results:** We propose to algorithmically edit gene expression data to drive cell-level morphological transitions using Generative Adversarial Networks (GAN) and GAN Inversion models. Leveraging cutting-edge spatial transcriptomic datasets with subcellular in-situ resolution and matched high-content imaging data, we propose an in-silico approach to quantify, model and imitate pathological processes in real-life clinical tissue samples.

**Availability and implementation:** The code and video demo is accessible via https://github.com/CTPLab/In-silico-editing

## Introduction

Recent advances in generative artificial intelligence (AI) (Bermano et al., 2022; Croitoru et al., 2023) have received significant attention and have set new standards for producing high-quality visual content. Several approaches, powered by Generative Adversarial Nets (GAN) and GAN Inversion (Xia et al., 2022), have been described to achieve remarkable editing effects on generated (Kang et al., 2023; Sauer et al., 2023) and real images (Patashnik et al., 2021). However, these methods mostly leverage encoded textual descriptions for desired image alterations. Typically, a pre-trained text encoder (Radford et al., 2021) is used to generate textual representations, which are then fed into the GAN (Inversion) model to guide the editing process.

In application to biomedicine, leveraging paired datasets of high-content biomedical imaging with multi-omic data holds great promise for in silico modelling of disease states and determining the effects of biologically targeted therapies. Engineering virtual cellular environments could aid the understanding of complex biological systems and enable low-cost and high-throughput morphological simulations. Previous research (Wu and Koelzer, 2023) has shown the feasibility of such simulations for cell-based drug screening. Here, we develop a novel integrative model of gene expression information and proteomics with morphological characteristics, and apply GAN (Inversion) models to model high-content cellular imaging with associated spatially resolved transcriptomic and proteomic data. Considering spatial transcriptomic (ST) datasets (Moses and Pachter, 2022) as multi-modal data instances, including high-plex mRNA transcripts as surrogates for gene expression and multi-channel histological images as phenotype representation, we propose to algorithmically edit the spatially localized expression of genes to drive morphological transitions.

## Materials and methods

In the experiments, we utilized the publically available CosMx (He et al., 2022) human liver and Xenium (Janesick et al., 2022) human lung datasets. The CosMx platform provides two comprehensive spatial expression maps of 1000-plex gene expression (approx. 0.2 billion gene expression counts for the normal liver slide and 0.5 billion for the hepatocellular carcinoma (HCC) slide), which correspond to 340k and 464k cells respectively^1^. Similarly, the Xenium dataset offers two large-scale spatial expression maps of 392-plex predesigned and custom target genes (approx. 24 million and 67 million total counts for the healthy lung slide and invasive adenocarcinoma (IAC) slide individually), along with 300k and 530k cells detected from the two slides ^2^.

### CosMx

As the protein biomarker CD298/B2M measured by multiplexed fluorescent imaging demonstrated strikingly different expression levels between the normal and tumor slides, we carefully confirmed the plausibility of protein expression patterns by comparing the image brightness in the fluorescent channel and gene expression levels of *HLA-A* and *B2M* (Fig. 3 in the Appendix), and therefore used this channel for the follow-up analysis. In combination with the DAPI channel, this forms two-channel images that are fed into the model training. Based on the clustering annotation provided in this dataset ^1^, we select normal cells (Hep 1, 3, 4, 5, 6 in Fig. 1 (f, left)) and hepatocellular carcinoma cells (Tumor 1 in Fig. 1 (f, right)) for an in-depth investigation of morpho-molecular changes between tumor and normal tissue.

**Fig. 1:**
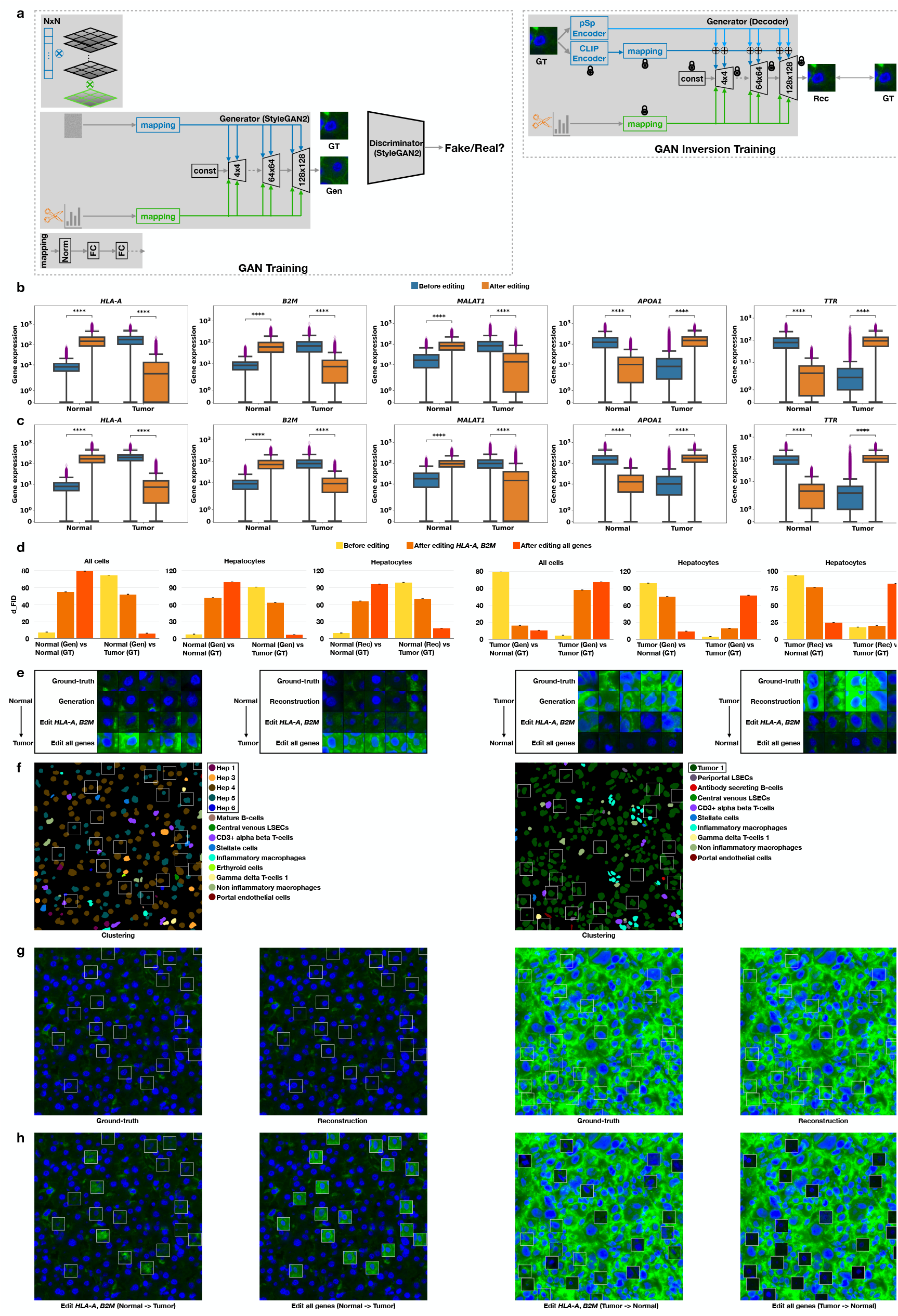
The model illustrations and experimental results for CosMx human liver dataset. **a**. The conceptual illustrations for the customized GAN and GAN Inversion model. **b**. The quantitative comparison of gene editing effects on all cells of the normal and tumor slide. Here, **** means *p* ≤ 0.0001. **c**. The quantitative comparison of gene editing effects on the ‘Hepatocyte’ subpopulation of the normal and tumor slide. **d**. The *d*_FID_ comparison of editing-driven cellular morphology transitions of the total and ‘Hepatocyte’ population for the generated cells (Gen) from the GAN training and the reconstructed cells (Rec) from the GAN Inversion training. Here, we randomly repeat the *d*_FID_ computation four times and report the mean and standard deviation. **e**. The image gallery of cellular morphological transitions of generated and reconstructed hepatocytes with reference to the ground-truth (GT) hepatocytes. Here, we present the transition that occurred in the DAPI (blue) and CD298/B2M (green) channels. **f**. The visualization of cell subtyping on a region of interest extracted from the normal and tumor slide. In the experiments, we focus on the cell subtypes “normal hepatocytes” (left plot, **Hep 1, 3, 4, 5, 6**) and “tumor hepatocytes” (right plot, **Tumor 1**). **g**. The randomly sampled ground-truth and reconstructed cellular images within the bounding boxes. **h**. The morphological transitions of these cellular images driven by targeted genes and all the genes.

### Xenium

In the Xenium IAC dataset we focus on normal lung epithelial cells and adenocarcinoma (tumor cells) as the populations of interest. We note that well-defined cell-level annotations are not available in the raw dataset^2^. We thus performed a careful evaluation of the histological images and cell-level clustering in the context of the expression of epithelial-related genes (See Fig. 4 in the Appendix), and found the second cluster of the ‘kmeans_2_clusters’ category supplied in the raw data to be highly enriched for cells of epithelial lineage for both the normal and tumor slide. Because both slides are single-channel DAPI images, we utilized the gray-scale cell-level images as the input for the Xenium experiment. Results of the Xenium analysis are presented in Fig. 2 of the Appendix.

**Fig. 2:**
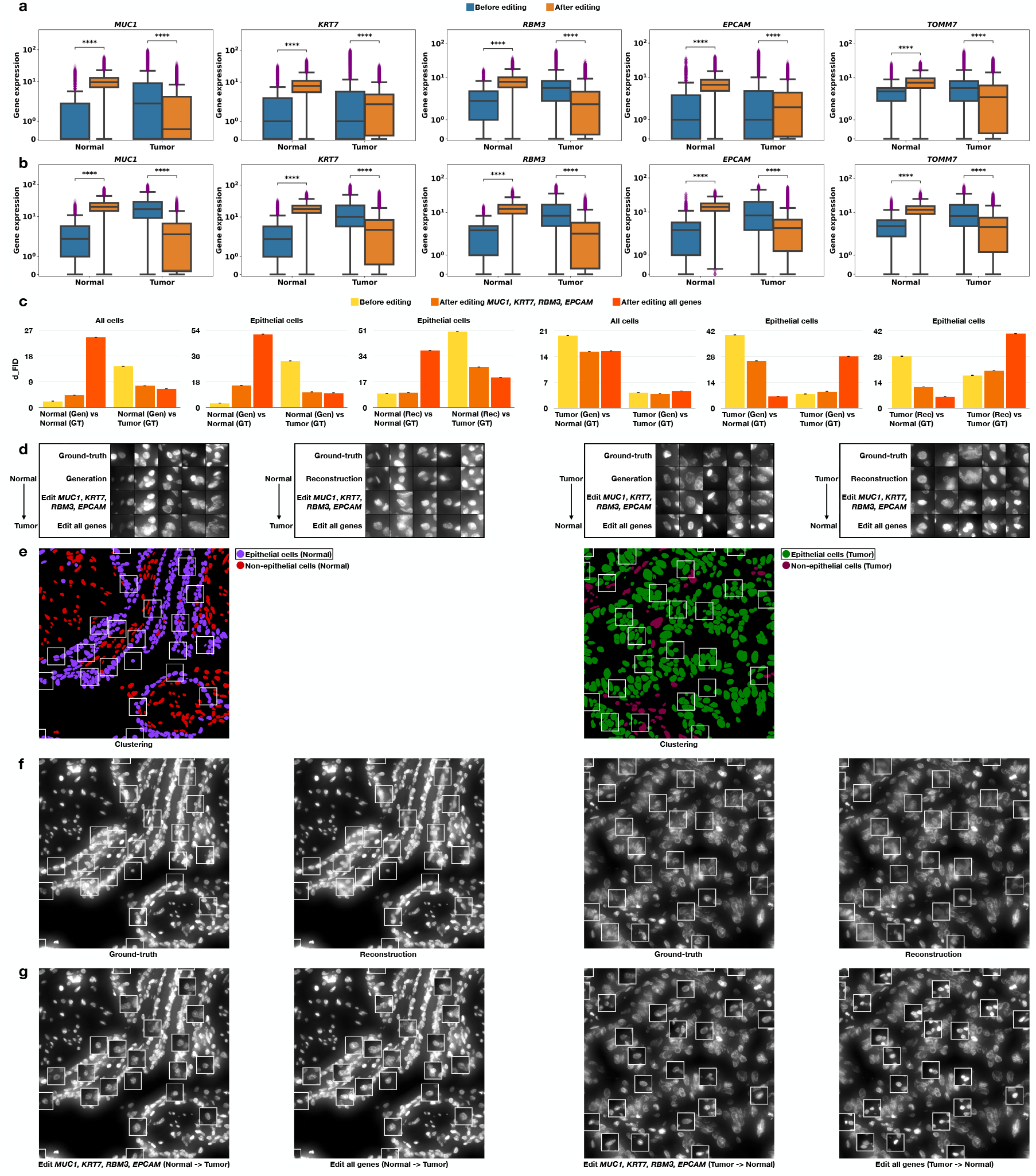
The experimental results for Xenium human lung dataset. **a**. The quantitative comparison of gene editing effects on all cells of the normal and tumor slide. Here, **** means *p* ≤ 0.0001. **b**. The quantitative comparison of gene editing effects on the ‘Epithelial cell’ subpopulation of the normal and tumor slide. **c**. The *d*_FID_ comparison of editing-driven cellular morphology transitions of the total and ‘Epithelial cell’ population for the generated cells (Gen) from the GAN training and the reconstructed cells (Rec) from the GAN Inversion training. Here, we randomly repeat the *d*_FID_ computation four times and report the mean and standard deviation. **d**. The image gallery of cellular morphology transitions of generated and reconstructed epithelial cells with reference to the ground-truth (GT) epithelial cells. Here, we present the transition that occurred in the DAPI channel. **e**. The cell subtyping visualization on a region of interest from the normal and tumor lung slide. In the experiments, we focus on the cell subtype of normal epithelial lung cells (left plot, **Epithelial cells (Normal)**) and adenocarcinoma cells (right plot, **Epithelial cells (Tumor)**) **f**. The randomly sampled ground-truth and reconstructed cellular images within the bounding boxes. **g**. The morphological transitions of these cellular images driven by targeted genes and all genes.

**Fig. 3:**
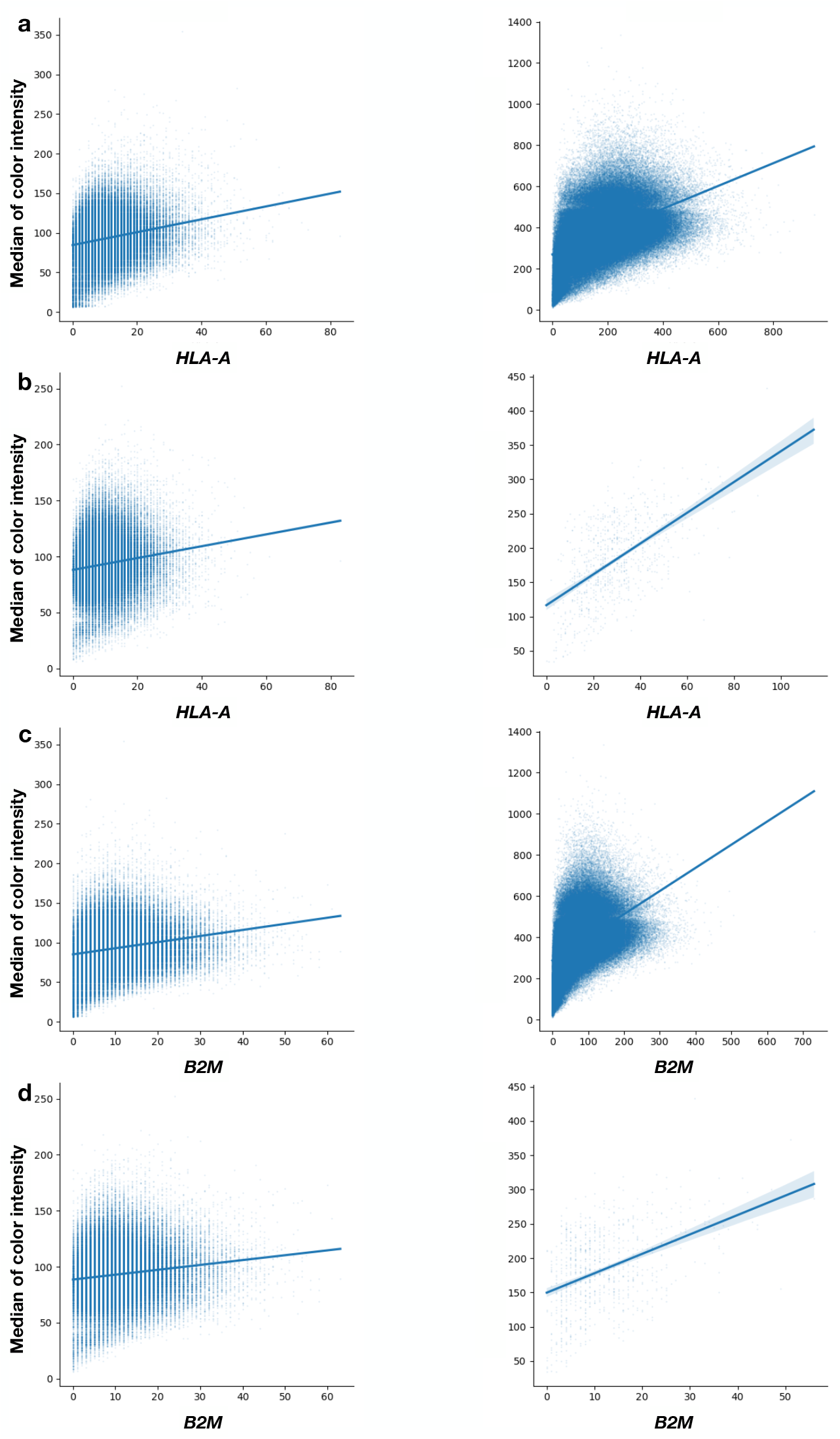
The plots of positively correlated median color intensity of CD298*/*B2M fluorescent marker and *HLA-A, B2M* gene expression for the CosMx human liver dataset. **a**. The plots of median color intensity of CD298/B2M and *HLA-A* for all cells from the normal (left) and tumor (right) slide. **b**. The plots of median color intensity of CD298/B2M and *HLA-A* for non-malignant hepatocytes from the normal (left) and tumor (right) slide. **c**. The plots of median color intensity of CD298/B2M and *B2M* for all cells from the normal (left) and tumor (right) slide. **d**. The plots of median color intensity of CD298/B2M and *B2M* for non-malignant hepatocytes from the normal (left) and tumor (right) slide.

**Fig. 4:**
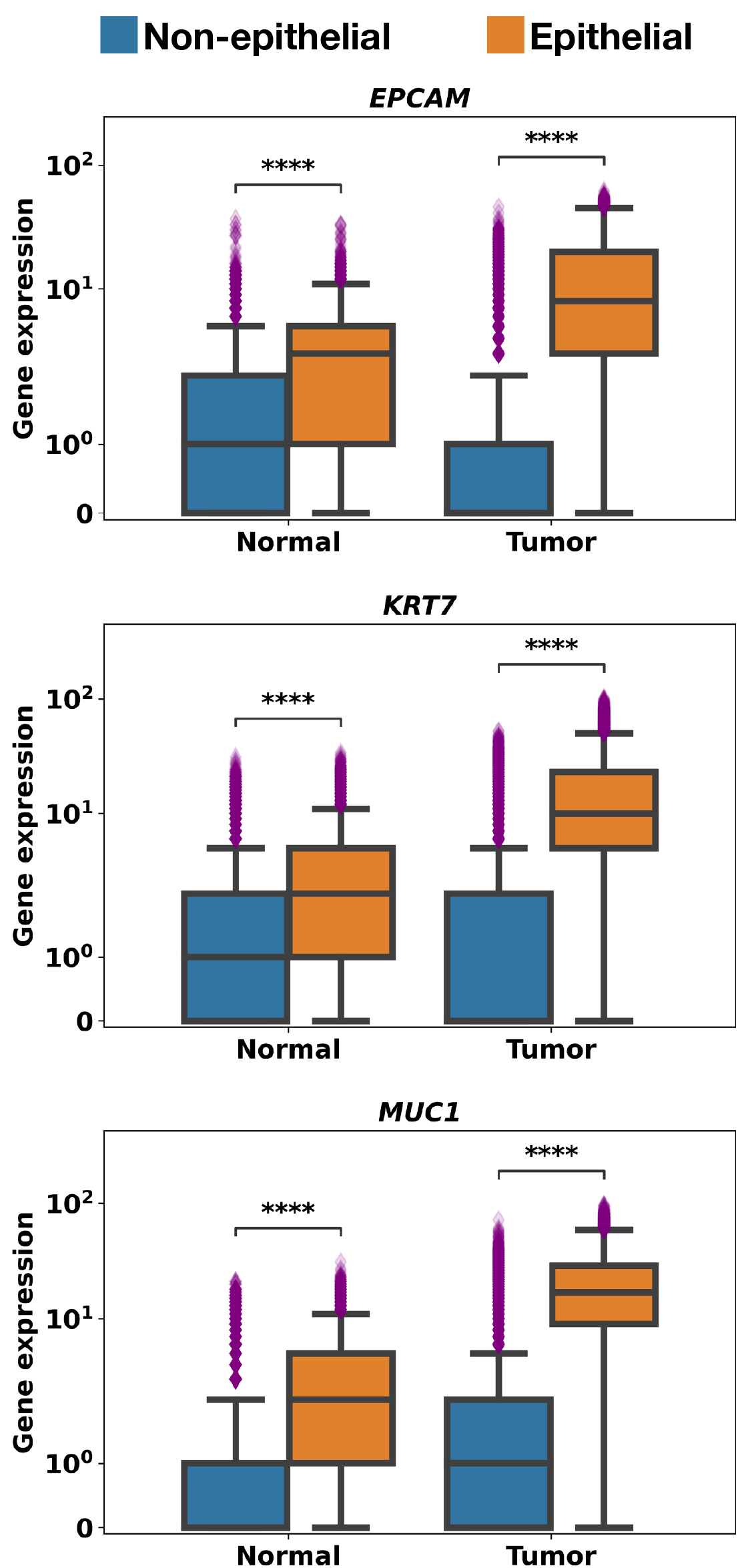
The quantitative comparison of the expression level of key genes of interest (*EPCAM* (top), *KRT7* (middle), *MUC1* (bottom)) that stratify the non-epithelial and epithelial-like lung cells of both the normal and tumor slide for the Xenium human lung dataset. Here, **** means p ≤ 0.0001.

### In silico gene expression editing

Customized on the gold-standard StyleGAN2 (Inversion) models (Karras et al., 2020; Alaluf et al., 2021) (Fig. 1 (a) and Appendix), we carry out_125_ interpretable morphological transitions of generated and real (reconstructed)_126_ cellular images by algorithmically editing the associated gene expression_127_ levels. Due to the lack of one-on-one correspondence between cells in the_128_ normal and tumor slides, clear guidance for individually editing the gene expression of each cell is missing. Instead, we propose to collectively edit the gene expression of cells by matching the gene data distribution (*i*.*e*., sample covariance matrix (SCM) (Wu and Koelzer, 2022)) of one cell population to another. This is done by scaling the eigenvalues and rotating the eigenbasis of SCM. For *i* = 0, 1 consider the 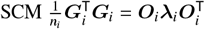, where ***G***_*i*_ is the collection of *n*_*i*_ *p*-plex gene expression from the normal or tumor slide, ***O***_*i*_ is the *p* × *p* eigenbasis and ***λ***_*i*_ is the *p* × *p* (sorted) diagonal eigenvalues obtained by eigenvalue decomposition. For the collection of ***G***_*i*_, we apply the linear transformation 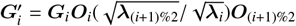 such that for the edited gene collection 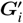 it holds 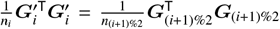. Due to the computational fluctuation of smaller eigenvalues and the dominant effect of the leading one, it suffices to scale the largest eigenvalue in our experiments. By keeping the selected group of gene expression unchanged during the linear transformation, we narrow down the editing process to the gene expression of interest. As shown in Fig. 1 (b, c) and Fig. 2 (a, b), the gene expression of interest has been clearly shifted towards the counterpart. After feeding the edited gene expression and corresponding center-cropped cellular image to the StyleGAN2 and StyleGAN2 Inversion model, we quantify and simulate the morphology transition at single-cell resolution.

## Results

First, we benchmark the model performance on all the cells for each experimental setting (Hepatocellular Carcinoma: plot 1 and 4 in 1 (d);_153_ Lung IAC: plot 1 and 4 in Fig. 2 (c)) from each slide and then narrow down to the critical cell subpopulations to derive biologically meaningful interpretations: (1) Normal hepatocytes versus tumor hepatocytes (HCC) for the CosMx dataset (Plots titled ‘Hepatocytes’ of Fig. 1 (d)); (2) Normal versus adenocarcinoma (tumor) epithelial cells for the Xenium IAC dataset (Plots titled ‘Epithelial cells’ of Fig. 2 (c)). We then investigate the editing effects of the top differentially expressed gene candidates with relevant biological functions as well as global gene expression patterns between normal and cancer tissues.

As a whole, we perform three levels of investigation that correspond to 8 variants. 1) Cell-level: Generated (StyleGAN2) and real reconstructed (StyleGAN2 Inversion) cellular images; 2) Gene expression-level: Expression of selected genes before and after in silico editing; 3) Population-level: The total cell population and clinically relevant cell subpopulations (‘Hepatocytes’ for CosMx and ‘Epithelial cells’ for Xenium).

At baseline, we witness a meaningful change in the expression of edited genes when comparing paired populations. As shown in Fig. 1 (b, c), two groups of genes with the highest expression difference between normal hepatocytes and HCC can be identified from the CoxMx dataset, *e*.*g*., genes with low expression in normal and upregulation in HCC (*HLA-A, B2M, MALAT1*) and genes downregulated in HCC compared to normal (*APOA1, TTR*). Following the in silico editing of normal cell types to the HCC space, the quantitative comparison presents a highly significant shift in gene expression levels towards the cancer spectrum for the total cell population (Fig. 1 (b)) and hepatocyte subpopulation (Fig. 1 (c), ∼80% of the total cells). Similar directional changes of expression levels can be observed for the genes of interest *MUC1, KRT7, RBM3, EPCAM, TOMM7* in the Xenium IAC dataset, though a clearer pattern of gene expression shift is demonstrated for the ‘Epithelial cell’ population (Fig. 2 (b), ∼20% of the total cells) than the total cell population (Fig. 2 (a), including diverse and heterogenous cell subtypes). After inputting edited genes of interest to the GAN and GAN Inversion model, we quantify the changes in cellular morphology using Fréchet Inception Distance (*d*_FID_) (Heusel et al., 2017), which effectively measures the statistical distance between compared image collections for the total cell population and the cell subtypes of interest.

As reported in Fig. 1 (d), after decreasing the leading and highly overexpressed genes *HLA-A* and *B2M* for the Cosmx HCC dataset, we achieve the quantifiable reversal of morphological features of tumor de-differentiation towards normal cells (decreasing nuclear size, decreased variation of nuclear size and shape). Further, more striking morphological effects become evident when editing the expression of all genes, in terms of clearly decreasing *d*_FID_ scores along both directions. Complementary to these quantification results, normal liver cells demonstrate a remarkably malignant appearance Fig. 1 (e, h) driven by edited genes of interest (increase in nuclear size, pronounced variation in nuclear size and shape). More importantly, the in silico editing of *HLA-A* and *B2M* indeed correlates to the emergence or disappearance of CD298/B2M protein expression as captured by the fluorescent imaging (green channel), supporting the reliability of biological interpretations brought by our in silico editing approach.

When analyzing editing effects of the Xenium lung dataset, we first observe the morphology-level heterogeneity for the total cell population, similar to the gene-level heterogeneity discussed above. However, expected trends of *d*_FID_ scores immediately emerge for the edited epithelial subtypes (Fig. 2 (c)), captured by the decreasing *d*_FID_ between transformed normal epithelial cell population and cancer cells. When narrowing down to the leading heterogeneous genes *MUC1, KRT7, RBM3* and *EPCAM*, the editing-driven cellular transition has proven to be effective *w*.*r*.*t*. the decreasing *d*_FID_ between the transformed normal cellular images and the tumor counterparts. As illustrated in the single-cell gallery of Fig. 2 (d) and cellular images within the tissue context of Fig. 2 (g), we demonstrate the emergence of atypical cellular features for normal epithelial liver cells driven by transforming the expression level of these four leading genes to the malignant spectrum.

## Conclusions

In conclusion, we verify the feasibility of the proposed method on two cutting-edge ST datasets. After algorithmically editing the expression of selected genes of the tumor cell population to match the expression level of matched normal pairs, we achieved quantifiable morphological transitions of cell-level images, indicating the effective reversal of tumor cell morphology to the healthy spectrum (and vice versa). The in silico editing approach, which exhibits low ethical, legal, and regulatory risks in the simulated intervention of human biological material, thus provides a new perspective to quantify, model, and imitate pathological processes in real-life clinical tissue samples.

## Author contributions statement

J.W. and V.H.K. conceived the research idea. J.W. implemented the algorithm and carried out the experiments. J.W. and V.H.K. analyzed the results. J.W. and V.H.K. drafted the manuscript. V.H.K. supervised the project.

## Acknowledgements

This study is funded by core funding of the University of Zurich to the Computational and Translational Pathology Lab led by V.H.K. at the Department of Pathology and Molecular Pathology, University Hospital and University of Zurich.

## Appendix

Throughout the study, we apply the GAN-based models for carrying out our experiments. Among many peer methods, the StyleGAN2 (Karras et al., 2020) architecture was considered the state-of-art (Bermano et al., 2022) and its key components have been continuously reused in scaled descendants (Sauer et al., 2023; Kang et al., 2023) that are competitive to the diffusion-based models (Croitoru et al., 2023). Therefore, we take StyleGAN2 as the backbone and customize it for our in silico editing on biomedical image data, where all the training experiments are run on an Nvidia DGX-A100 workstation. Given profound insights that can be derived from spatially resolved single-cell datasets (Stuart and Satija, 2019) and the technical feasibility that promises good reconstruction quality for cell-level images, we take pairs of center-cropped single-cell images with their associated gene expression vector as the ‘image’ and ‘text’ input modality of StyleGAN2 (Inversion) models. Based on the pathologist review and recommendation, we center-crop 160 × 160 and 96 × 96 resolution of the cellular image and associated sparse 1000/392-plex gene expression array for the CosMx and Xenium dataset using cell segmentation annotations provided in the raw data. Then, we uniformly re-scale the cropped images to 128 × 128 and form the paired one-dimensional tabular of gene expression by summing up the sparse array along each gene dimension for efficient GAN (Inversion) training.

### StyleGAN2

Because human genes directly encode the synthesis of proteins and other molecules that regulate the cellular structure, function, and behavior, we do not pre-train a large-scale text encoder to process gene expression. Instead, we propose to feed the gene expression vector to a simple mapping network of StyleGAN2 (Fig. 1 (a, left)) for regulating the morphological features. As a result, our biomedical interpretations are derived from gene expression levels rather than convoluted latent representations used in text-guided image generation. Aside from the standard adversarial loss, we also take gene expression as input of the class condition (*i*.*e*., cells from the normal (Class 0) or tumor slide (Class 1)) and project them to the last discriminator layer for class-based conditional GAN training. For both datasets, we set the batch size to 16 and train the customized StyleGAN2 model on the pair of center-cropped cellular images and their gene expression vectors for 800k iterations. Then, we freeze the best model determined by *d*_FID_ on the normal and tumor cell classes (Fig. 5 in the Appendix) and take it as the decoder for the GAN Inversion training. For *i* = 0, 1, consider ***μ***_*i*_, ***S***_*i*_ be the mean and covariance of the collection of cellular image representations of one slide, then we have 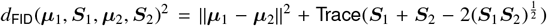, where we compute the *d*_FID_ with a more efficient implementation (Wu and Koelzer, 2022).

**Fig. 5:**
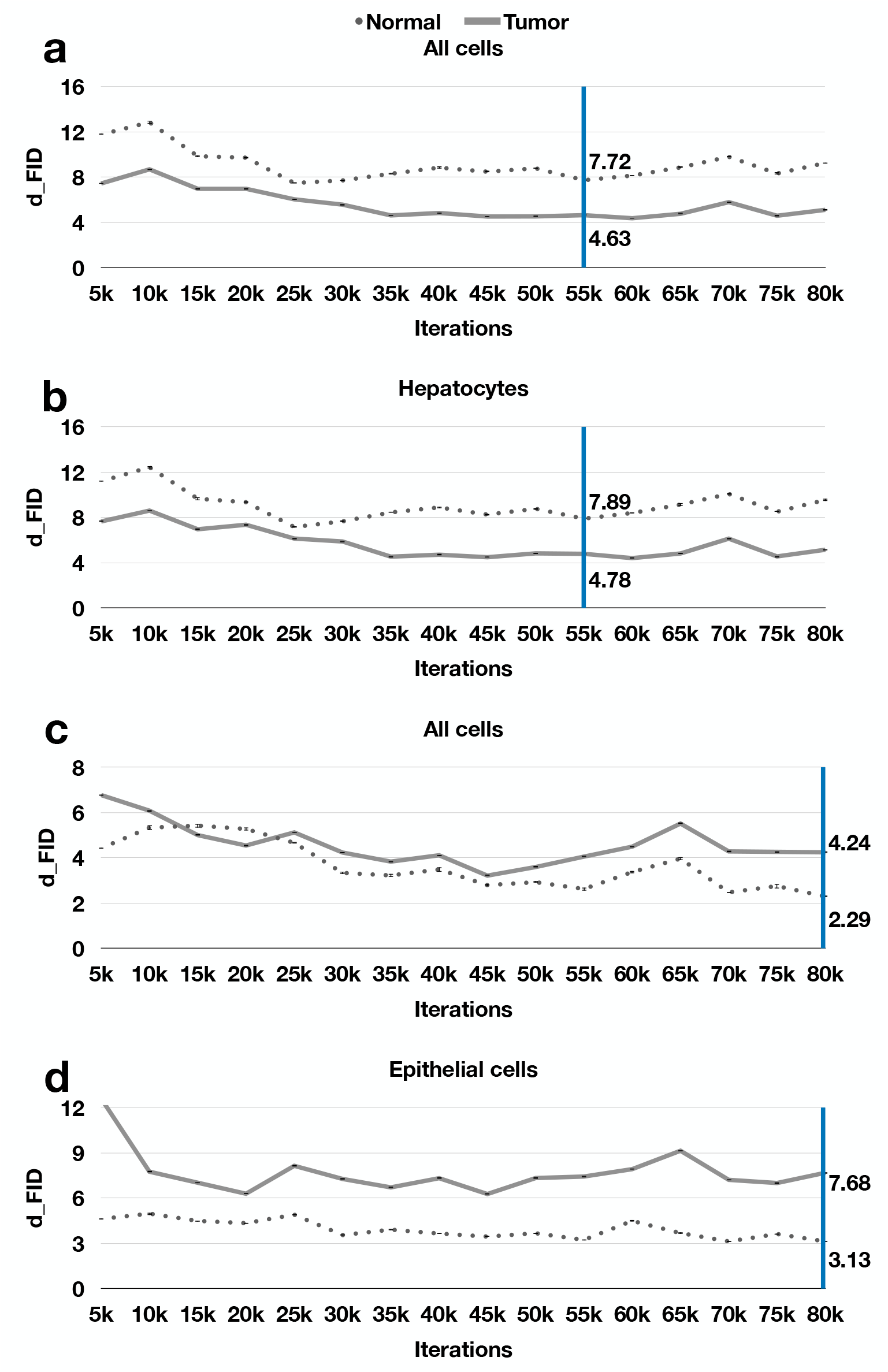
The *d*_FID_ results for GAN training. Here, we randomly repeat the *d*_FID_ computation four times and report the mean and standard deviation. The overall best *d*_FID_ scores are highlighted for each case. **a**. The *d*_FID_ curve for all cells from the normal and tumor liver slide of the CosMx dataset. **b**. The *d*_FID_ curve for all hepatocytes from the normal and tumor liver slide of the CosMx dataset. **c**. The *d*_FID_ curve for all cells from the normal and tumor lung slide of the Xenium dataset. **d**. The *d*_FID_ curve for all epithelial cells from the normal and tumor lung slide of the Xenium dataset.

### StyleGAN2 Inversion

Although meaningful interpretations can be derived from the generated (virtual) cellular images, we are also keen on verifying the editing effects on the real cells extracted from the two slides. This motivates us to utilize the GAN Inversion methodology (Xia et al., 2022) and analyze the morphological transitions that occur on real reconstructed (Rec) cellular images. To this end, we reuse the StyleGAN2 generator and freeze the model weights learned from the GAN training. As the dual goal of good reconstruction quality and editability presents an inherent challenge (Bermano et al., 2022), we additionally pre-train a simple ResNet-based CLIP encoder (Shariatnia, 2021) for image encoding and feed the output to the mapping function of our StyleGAN2 model for processing the noisy morphological representations (Fig. 1 (a, right)), which form the more editable part of representations for the downstream analysis. Meanwhile, we plug the ‘pixel2style2pixel’ (Richardson et al., 2021) (pSp) encoder that is proven to be effective in reconstructing high-fidelity natural images to the StyleGAN2 Inversion model and learn the latent morphological representations to achieve good reconstruction quality. During the training, we utilize the same objective suggested by Alaluf *et al*. (Alaluf et al., 2021) except the ID similarity loss used for human face images, *i*.*e*., ℒ = *γ*_1_ ℒ _moco_ + *γ*_2_ℒ _PIP_ + *γ*_3_ℒ _2_, where ℒ_moco_ is the contrastive loss, ℒ_PIP_ is the perceptually learned loss (Zhang et al., 2018), and ℒ_2_ is the *l*_2_ reconstruction loss. We set the batch size to 8 and train the inversion model for 800k iterations. Eventually, we perform the downstream analysis by the model with the overall best peak signal-to-noise Ratio (PSNR) and Structural Similarity Index Measure (SSIM) scores, as highlighted in Fig. 6 of the Appendix and image results of Fig. 1 (e, g) and Fig. 2 (d, f).

**Fig. 6:**
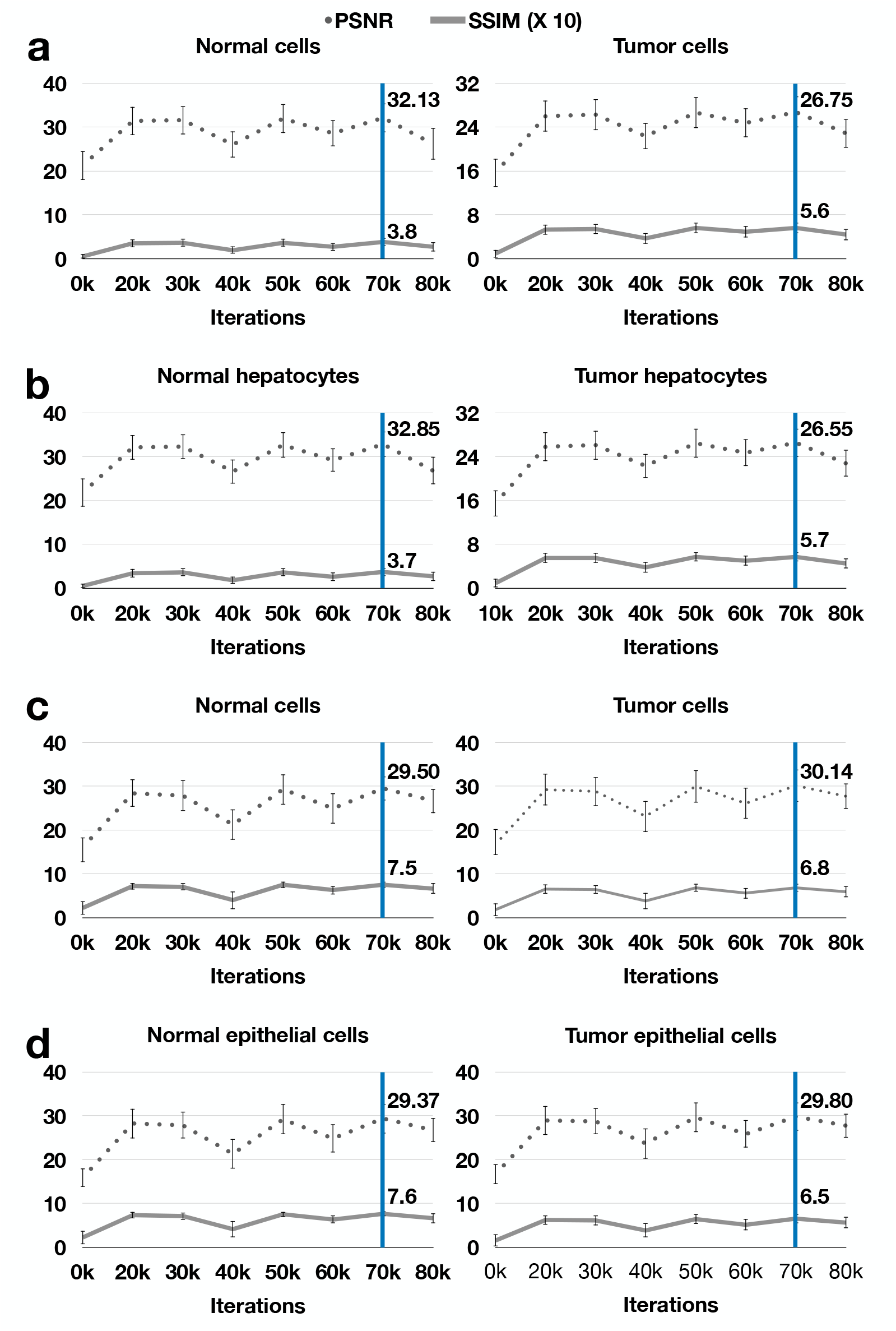
The PSNR and SSIM results for GAN Inversion training. Here, we report the mean and standard deviation for both measurements. The overall best PSNR and SSIM scores are highlighted for each case. **a**. The PSNR and SSIM curve for all cells from the normal and tumor liver slide of the CosMx dataset. **b**. The PSNR and SSIM curve for all hepatocytes from the normal and tumor liver slide of the CosMx dataset. **c**. The PSNR and SSIM curve for all cells from the normal and tumor lung slide of the Xenium dataset. **d**. The PSNR and SSIM curve for all epithelial cells from the normal and tumor lung slide of the Xenium dataset.

## Footnotes

1 https://nanostring.com/products/cosmx-spatial-molecular-imager/human-liver-rna-ffpe-dataset/

2 https://www.10xgenomics.com/resources/datasets/xenium-human-lung-preview-data-1-standard

## References

Y. Alaluf, O. Patashnik, and D. Cohen-Or. Restyle: A residual-based stylegan encoder via iterative refinement. In Proceedings of the IEEE/CVF International Conference on Computer Vision, pages 6711–6720, 2021.

A. H. Bermano, R. Gal, Y. Alaluf, R. Mokady, Y. Nitzan, O. Tov, O. Patashnik, and D. Cohen-Or. State-of-the-art in the architecture, methods and applications of stylegan. In Computer Graphics Forum, volume 41, pages 591–611. Wiley Online Library, 2022.

F.-A. Croitoru, V. Hondru, R. T. Ionescu, and M. Shah. Diffusion models in vision: A survey. IEEE Transactions on Pattern Analysis and Machine Intelligence, 2023.

S. He, R. Bhatt, C. Brown, E. A. Brown, D. L. Buhr, K. Chantranuvatana, P. Danaher, D. Dunaway, R. G. Garrison, G. Geiss, et al. High-plex imaging of rna and proteins at subcellular resolution in fixed tissue by spatial molecular imaging. Nature Biotechnology, 40(12):1794–1806, 2022.

M. Heusel, H. Ramsauer, T. Unterthiner, B. Nessler, and S. Hochreiter. Gans trained by a two time-scale update rule converge to a local nash equilibrium. Advances in neural information processing systems, 30, 2017.

A. Janesick, R. Shelansky, A. D. Gottscho, F. Wagner, M. Rouault, G. Beliakoff, M. F. de Oliveira, A. Kohlway, J. Abousoud, C. A. Morrison, et al. High resolution mapping of the breast cancer tumor microenvironment using integrated single cell, spatial and in situ analysis of ffpe tissue. bioRxiv, pages 2022–10, 2022.

M. Kang, J.-Y. Zhu, R. Zhang, J. Park, E. Shechtman, S. Paris, and T. Park. Scaling up gans for text-to-image synthesis. In Proceedings of the IEEE/CVF Conference on Computer Vision and Pattern Recognition, 2023.

T. Karras, S. Laine, M. Aittala, J. Hellsten, J. Lehtinen, and T. Aila. Analyzing and improving the image quality of stylegan. In Proceedings of the IEEE/CVF conference on computer vision and pattern recognition, pages 8110–8119, 2020.

L. Moses and L. Pachter. Museum of spatial transcriptomics. Nature Methods, 19(5):534–546, 2022.

O. Patashnik, Z. Wu, E. Shechtman, D. Cohen-Or, and D. Lischinski. Styleclip: Text-driven manipulation of stylegan imagery. In Proceedings of the IEEE/CVF International Conference on Computer Vision, pages 2085–2094, 2021.

A. Radford, J. W. Kim, C. Hallacy, A. Ramesh, G. Goh, S. Agarwal, G. Sastry, A. Askell, P. Mishkin, J. Clark, et al. Learning transferable visual models from natural language supervision. In International conference on machine learning, pages 8748–8763.PMLR, 2021.

E. Richardson, Y. Alaluf, O. Patashnik, Y. Nitzan, Y. Azar, S. Shapiro, and D. Cohen-Or. Encoding in style: a stylegan encoder for image-to-image translation. In Proceedings of the IEEE/CVF Conference on Computer Vision and Pattern Recognition, pages 2287–2296, 2021.

A. Sauer, T. Karras, S. Laine, A. Geiger, and T. Aila. Stylegan-t: Unlocking the power of gans for fast large-scale text-to-image synthesis. International Conference on Machine Learning, 2023.

M. M. Shariatnia. Simple CLIP, 4 2021.

T. Stuart and R. Satija. Integrative single-cell analysis. Nature reviews genetics, 20(5):257–272, 2019.

J. Wu and V. Koelzer. Sorted eigenvalue comparison d_Eig_: A simple alternative to d_FID_. In NeurIPS 2022 Workshop on Distribution Shifts: Connecting Methods and Applications, 2022.

J. Wu and V. H. Koelzer. Gilea: Gan inversion-enabled latent eigenvalue analysis for phenome profiling and editing. bioRxiv, pages 2023–02, 2023.

W. Xia, Y. Zhang, Y. Yang, J.-H. Xue, B. Zhou, and M.-H. Yang. Gan inversion: A survey. IEEE Transactions on Pattern Analysis and Machine Intelligence, 2022.

R. Zhang, P. Isola, A. A. Efros, E. Shechtman, and O. Wang. The unreasonable effectiveness of deep features as a perceptual metric. In Proceedings of the IEEE conference on computer vision and pattern recognition, pages 586–595, 2018.

